# Use of a ubiquitous gene-editing tool in budding yeast causes off-target repression of neighboring gene protein synthesis

**DOI:** 10.1101/2022.06.27.497784

**Authors:** Emily Nicole Powers, Lidia Llacsahuanga Allcca, Ella Doron-Mandel, Jenny Kim Kim, Marko Jovanovic, Gloria Ann Brar

## Abstract

Precision genome-editing approaches have long been available in budding yeast, enabling introduction of gene deletions, epitope tag fusions, and promoter swaps through a selection-based strategy. Such approaches allow loci to be modified without disruption of coding or regulatory sequences of neighboring genes. Use of this approach to delete *DBP1* however, led to silencing of expression and the resultant loss of function for the neighboring gene *MRP51*. We found that insertion of a resistance cassette to delete *DBP1*, drove a 5’ extended alternative transcript for *MRP51* which dampened Mrp51 protein synthesis. Misregulation of *MRP51* occurred through an integrated transcriptional and translational repressive long undecoded transcript isoform (LUTI)-based mechanism that was recently shown to naturally regulate gene expression in yeast and other organisms. Cassette-induced *MRP51* repression drove all mutant phenotypes we detected in cells deleted for *DBP1*. Selection cassette-mediated aberrant transcription events are not specific to this locus or a unique cassette but can be prevented by insertion of transcription insulators flanking the cassette. Our study suggests the existence of confounding off-target mutant phenotypes resulting from misregulated neighboring loci following genome edits in yeast. Furthermore, features of LUTI-based regulation are broadly conserved to eukaryotic organisms which indicates the potential that similar misregulation could be unnoticed in other edited organisms as well.

## Introduction

Genome engineering to create targeted gene deletions, mutations, and epitope-tagged proteins is a key strategy for the mechanistic dissection of almost any biological process. Budding yeast was the first eukaryotic organism in which this became highly facile, thanks to the development of a one-step PCR-based editing strategy, frequently used with a shared toolkit of selection cassettes^1–5^. These tools accelerated the use of yeast as a model organism and have become common, routinely used by thousands of labs, as well as enabling large global endeavors, like creation of the yeast deletion collection^6^. The utility of this approach stems from the ease of creating targeted mutations, simply by designing primers that amplify a cassette of choice while adding homologous sequences that flank the genomic region to be edited^1-5^. After a cassette is amplified, the linear PCR product is transformed into yeast and incorporated into the targeted site within the genome by cellular homologous recombination machinery^1–5^. Finally, selection for a cassette-encoded resistance marker allows one to isolate a clonal population of edited cells. Cassette incorporation is highly specific to the targeted genomic location and mutant strains created in this manner are typically regarded with high confidence^1–5^. Because of the ease of use and rapid nature of this selection-mediated editing strategy, it has remained commonplace despite the development of CRISPR/Cas9-based editing methods which allow the creation of “seamless” non-selection-marked mutations^7^.

While selection-mediated editing strategies have been used to advance countless discoveries in yeast, their utility relies on the assumption that insertion of a selection cassette at one locus will not disrupt the expression of neighboring loci. However, a growing body of evidence suggests that yeast genomic loci are not the simple discrete units they were long thought to be, but instead can have complex and linked transcriptional outputs. For example, it is now understood that activity from one transcription start site (TSS) can interfere with the output of nearby TSSs in an adjacent sense or antisense configuration, and that many promoters are bidirectionally active^8–19^. Transcription interference is seen in diverse organisms, including yeast and human cells, and can occur between TSSs driving coding or noncoding RNAs and controlling both transcript isoform identity and transcript levels^15,15,20–29^. It has also been shown that more than one TSS is often present at a single locus, even in the simple budding yeast^14^.

Transcription interference between TSSs within the same genomic locus is an important feature of a naturally occurring type of regulation dependent on LUTIs (long undecoded transcript isoforms)^20,30^. For genes regulated by this strategy, 5’ extended LUTIs are transcribed in place of canonical mRNAs to temporally downregulate protein synthesis^20,30–32^. Here, use of an upstream alternate TSS represses transcription from the canonical TSS in cis-via transcriptional interference^20,30^. In concert, translation of the main ORF-encoded protein products from LUTIs are repressed by translation of competitive AUG-initiated upstream ORFs (uORFs)^30,33–37^. Natural LUTI-based regulation has been shown to modulate expression for many genes during meiosis and the unfolded protein response in yeast^30–32^. Features of LUTI-based regulation are conserved across eukaryotes, and evidence suggests this regulation may also occur in more complex eukaryotes such as during mammalian embryonic stem cell differentiation and larval development in *Drosophila melanogaster*^38,39^. Interestingly, we find that insertion of resistance cassettes commonly used to mark genomic edits in yeast, can induce the repression of neighboring genes though synthetic and constitutive LUTI-based regulation.

We report that standard selection-cassette replacement of the *DBP1* ORF causes mis-regulation of the adjacent gene *MRP51*, by aberrant LUTI-based repression, highlighting an overlooked side-effect of this genome engineering strategy. Every phenotype we observed in cells with a cassette-mediated deletion of the *DBP1* ORF was caused by cassette-induced mis-regulation of the neighboring gene *MRP51*, rather than loss of *DBP1*. Non-specific cassette-induced phenotypes were general to all selection cassettes that we tested to replace the *DBP1* ORF. These data suggest the standard method for making selection-marked mutations in yeast may drive unexpected side effects that have thus far been overlooked. Consistent with this, we found that cassette driven aberrant transcripts are not unique to the *DBP1* locus and can be detected in roughly 25% of loci edited in this manner. While this situation may suggest the use of “seamless” genome editing methods to avoid cassette-driven artifacts, we find that even CRISPR/Cas9 unmarked mutations can introduce surprising effects on regulation of mutated loci. This, as well as the increased ease of use of selection-marked mutations, motivated us to design and test a panel of “insulated” cassettes that prevent aberrant transcription from interfering with neighboring genes *in vivo*. Use of these cassettes in yeast should improve the specificity of engineered mutant strains for future studies. More broadly, our study highlights a long-overlooked and potent side effect that can result from genome-editing, and that may impact published studies performed both in yeast and more complex eukaryotes.

## Results

### Insertion of a resistance cassette at the *DBP1* locus causes aberrant transcription and reduced protein production from the neighboring gene *MRP51*

We had previously observed upregulation of the Dbp1 helicase in meiotic yeast cells, coincident with downregulation of its paralog, Ded1^40^. Ded1 is important for translation initiation, and has been well studied under conditions of mitotic exponential growth^41,42^. In contrast, the role of Dbp1 has remained less clear, partially a result of its absent or low expression under commonly studied laboratory conditions. Based on the identity of Dbp1 as a RNA helicase^43^, we predicted that Dbp1 could have a role in regulating gene expression during meiosis. To test this, we deleted *DBP1* by replacing the ORF (from start to stop codon) with a standard cassette encoding Geneticin (G418)-resistance, amplified from the pFA6a-KANMX6 plasmid^1^, in the SK1 budding yeast strain background^44^. We then compared gene expression profiles of wild-type and *dbp1Δ::KANMX6* cells during meiosis, using ribosome profiling and mRNA-sequencing (mRNA-seq). To our surprise, *MRP51*, the gene located directly adjacent to *DBP1*, exhibited a profound decrease in translation in the *dbp1Δ::KANMX6* strain relative to wild-type controls (Figure 1A). Translation of *MRP51* was 8.4-fold lower in *dbp1Δ::KANMX6* cells despite a 1.8-fold increase in *MRP51* mRNA abundance (Figure 1A-B). These changes reflected a 15-fold decrease in *MRP51* translation efficiency (TE) in the *dbp1Δ::KANMX6* cells compared to the wild-type control (Figure 1C; TE: ribosome footprint FP RPKM/ mRNA RPKM) and led to a ∼40% reduction in Mrp51 protein level compared to wild type, as assessed by western blot (Figure 1D). Because of the close genomic proximity of these genes, we hypothesized that this change in TE could be due to cis effects from the *dbp1Δ::KANMX6* insertion rather than regulation of *MRP51* by Dbp1 protein. Upon closer examination of the mRNA transcripts produced at this locus, we found that replacing the *DBP1* ORF with the KANMX6 resistance cassette led to production of a 5’ extended *MRP51* mRNA compared to that in wild-type cells. The extended transcript contained 3 AUG-initiated upstream ORFs (uORFs) not present on the wild-type transcript, which were translated at the expense of the *MRP51* ORF in *dbp1Δ::KANMX6* cells (Figure 1E).

**Figure 1.**
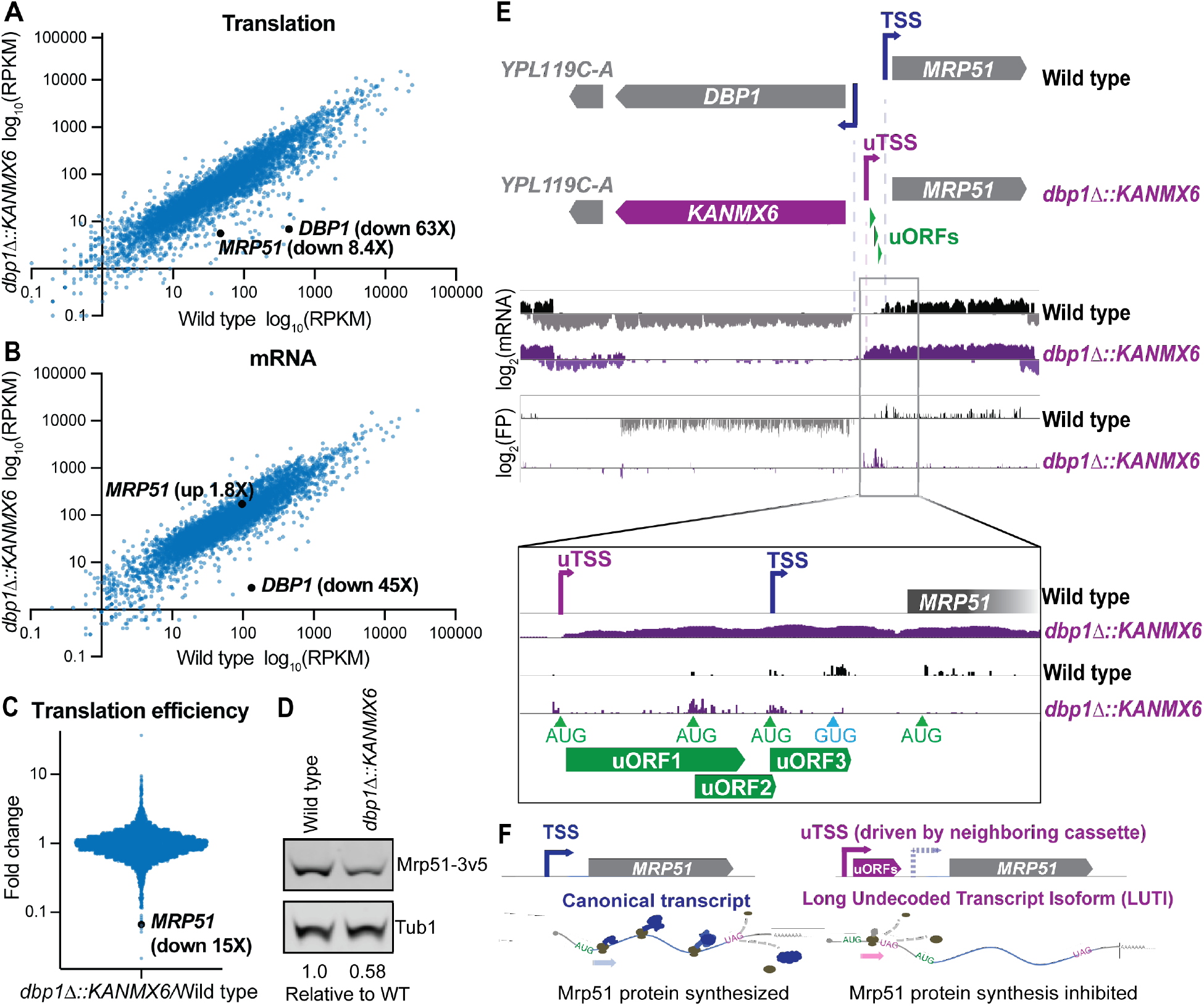
Insertion of a resistance cassette at the *DBP1* locus causes aberrant transcription and reduced protein production from the neighboring *MRP51* gene. (A) Translation (ribosome profiling FP) and (B) mRNA-seq reads per kilobase million mapped reads (RPKM) for every ORF expressed in wild-type and *dbp1Δ::KANMX6* cells. (C) Fold-change of translation efficiency (TE: FP RPKM/mRNA RPKM) for all expressed genes. For RPKM for all quantified genes, see Figure 1 – Source Data 1. (D) Western blots of mitotic yeast demonstrate that Mrp51 protein level is decreased to 58% that of wild type in *dbp1Δ::KANMX6* cells. Mrp51 levels were normalized to alpha tubulin. For full blot scans see Figure 1 – Source Data 1. (E) mRNA and FP reads mapped to the *DBP1/MRP51* loci in wild-type (grey/black) and *dbp1Δ::KANMX6* (purple) cells. Note that *MRP51* encoding transcripts are 5’ extended in the *dbp1Δ::KANMX6* cells compared to wild type (see mRNA tracks). The extended transcript contains 3 AUG initiated uORFs translated at the expense of the *MRP51* ORF (see FP tracks and inset). (F) Model: replacement of *DBP1* ORF with a resistance cassette causes aberrant expression of a long undecoded transcript isoform (LUTI) for *MRP51*, which results in lower Mrp51 protein expression as an off-target effect.

These data suggest that KANMX6 cassette insertion at the *DBP1* locus drives the transcription of an aberrant 5’ extended *MRP51* mRNA containing repressive uORFs and causing reduced Mrp51 protein production. Thus, the features of *MRP51* mis-regulation in this strain, mimicked the natural LUTI-based mode of regulation that conditionally downregulates protein synthesis from many genes in yeast (Figure 1F)^20,30–32^. Here, the primary difference was that repression of *MRP51* expression appears to occur constitutively as result of KANMX6 insertion at the neighboring *DBP1* ORF rather than as a result of temporally regulated toggling between two TSSs.

Because of the widespread and long-term use of cassette-mediated gene deletion in yeast, we were surprised to observe this dramatic off-target effect. To confirm aberrant *MRP51* regulation was not an artifact of our strain, experimental conditions, or selection cassette, we analyzed a published ribosome profiling and mRNA-seq dataset comparing wild type and *dbp1Δ* cells of the S288C budding yeast background. These *dbp1Δ* cells were generated by replacement of the *DBP1* ORF with a cassette encoding resistance to Hygromycin (*dbp1Δ::HYGMX4*)^45^. Consistent with our data, insertion of HYGMX4 to replace the *DBP1* ORF caused mis-regulation of the adjacent gene *MRP51*. Despite levels of the Mrp51-encoding transcript remaining similar to wild type in *dbp1Δ::HYGMX4* cells, translation of *MRP51* was decreased 4.4-fold in these conditions (Figure 1—figure supplement 1A-B). The 3.7-fold decrease in TE of *MRP51* in *dbp1Δ::HYGMX4* cells in this study led to the interpretation that the canonical *MRP51* transcript is highly dependent on Dbp1 for its translation (Figure 1—figure supplement 1C). A closer look at the transcripts produced from this locus, however, revealed the presence of a 5’-extended *MRP51* LUTI in the *dbp1Δ::HYGMX4* cells, like the one we had observed. This 5’ extended *MRP51* transcript contained competitive AUG-initiated uORFs that were translated in place of the *MRP51* ORF, as in our experiments (Figure 1—figure supplement 1D). Together these data indicate that cassette replacement of the *DBP1* ORF drives aberrant transcription independent of strain, cassette selection identity, and experimental conditions. Instead, the observed mis-regulation of *MRP51* appears to be a general side-effect of the cassette-insertion.

**Figure 1—figure supplement 1.**
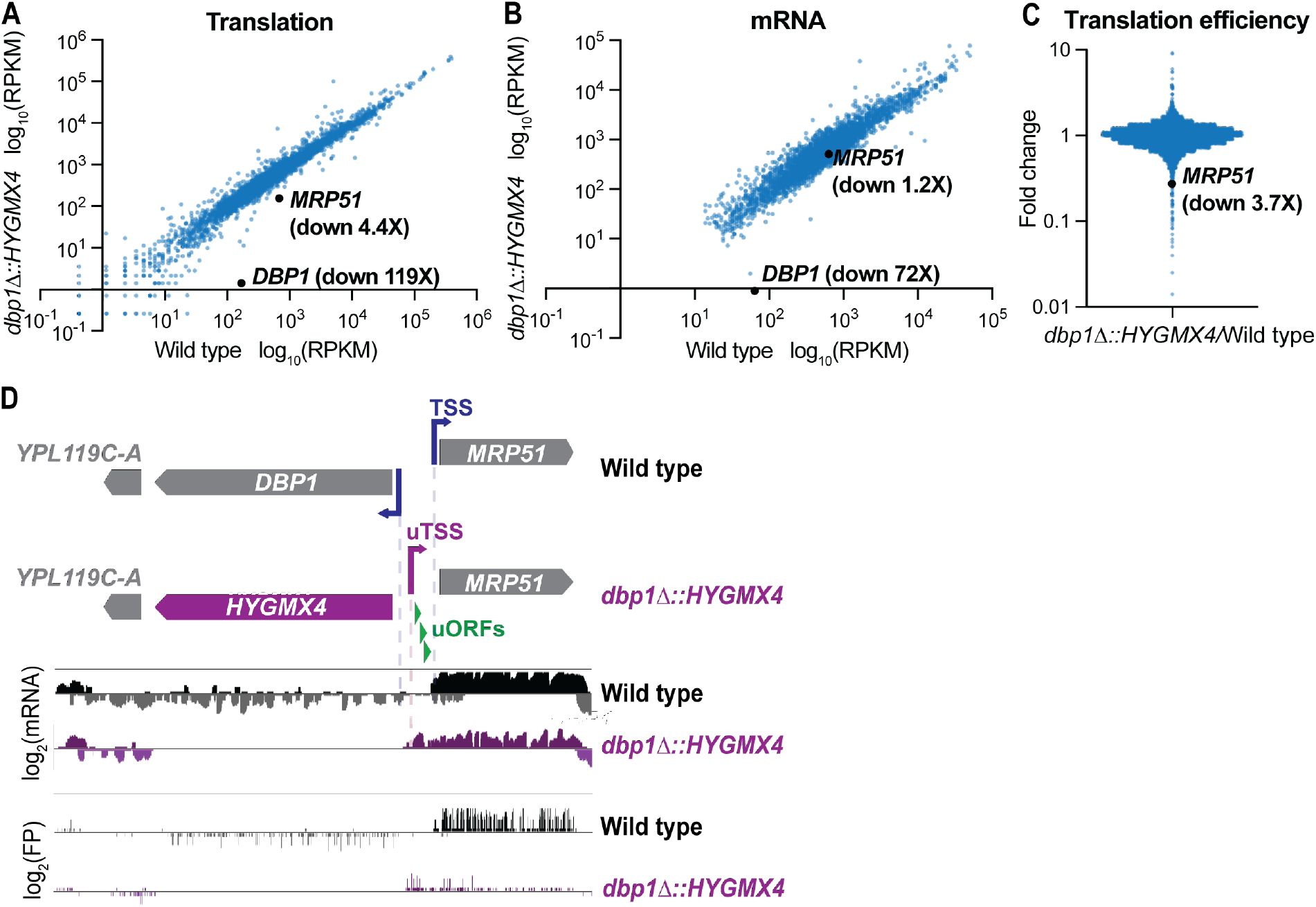
A published dataset confirms aberrant transcription and mis-regulation of *MRP51* following cassette-mediated *DBP1* replacement. Ribosome profiling and mRNA-seq of *dbp1Δ::HYGMX4 and* wild-type S288C budding yeast during mitotic growth from Sen et al., 2019. (A) Translation (FP RPKM) and (B) mRNA RPKM counts of all genes expressed. (C) Fold-change TE for every gene expressed in *dbp1Δ::HYGMX4* and wild-type cells. (D) mRNA and FP reads aligned to the *DBP1/MRP51* loci for wild type (grey/black) and *dbp1Δ::HYGMX4* (purple). Note the *MRP51* mRNA is 5’ extended in the *dbp1Δ::HYGMX4* cells (mRNA tracks) and the translation of uORFs not present on the wild-type mRNA (FP tracks).

### Mis-regulation of *MRP51* by insertion of a resistance cassette leads to systemic phenotypic consequences

We next sought to determine whether the mutant phenotypes that we observed in *dbp1Δ::KANMX6* cells were due to loss of *DBP1* function or resistance cassette-dependent mis-regulation of *MRP51*, a nuclear encoded mitochondrial small subunit ribosome protein (mt-SSU)^46^. Polysome analysis of *dbp1Δ::KANMX6* cells during meiosis demonstrated that overall translation rates were diminished relative to wild-type cells, made evident by lower polysome peaks in the mutant (Figure 2A; right, fractions 5-6). Consistent with this finding, measurement of radioactive amino acid incorporation rates revealed a 24% decrease in bulk translation in *dbp1Δ::KANMX6* cells (Figure 2A—figure supplement 1A). Polysome traces also indicated differences in the accumulation of a ribonucleoprotein (RNP) species roughly the size of the cytoplasmic large 60S subunit (Figure 2A; LSU, fraction 3) and a decrease in cytoplasmic small 40S subunit signal (Figure 2A; SSU, fraction 2). We next performed label-free mass spectrometry on fractionated wild-type and *dbp1Δ::KANMX6* polysomes and used hierarchical clustering of the data to assess global differences in polysome composition. To our surprise, we identified a prominent cluster of proteins enriched in fraction 3 of polysomes from *dbp1Δ::KANMX6* cells compared to wild type, that was highly enriched for the mitochondrial large ribosome subunit (mt-LSU). This cluster contained 34 of the 42 proteinaceous mt-LSU components quantified in our study (Figure 2A; left). Further analysis of fraction 3 revealed every mitochondrial 54S protein component quantified in this fraction was enriched in *dbp1Δ::KANMX6* cells compared to the wild-type control (Figure 2—figure supplement 1B). These data suggest the accumulated RNP observed in *dbp1Δ::KANMX6* cells is the mt-LSU. A previous study from our lab observed that when individual cytoplasmic SSU proteins are lost, free cytoplasmic 60S LSUs accumulate, presumably a result of their inability to find a SSU to complex with and form fully assembled 80S species.^47^ The build-up of free mt-LSU in *dbp1Δ::KANMX6* cells, which express lower amounts of mitochondrial small subunit (mt-SSU) component Mrp51 is reminiscent of that effect. Furthermore, we identified a small cluster of cytoplasmically translated mitochondrial proteins that were decreased in *dbp1Δ::KANMX6* high polysome fractions compared to wild type, hinting at effects of broader mitochondrial dysfunction. These data suggested the cassette-mediated mis-regulation of *MRP51* in *dbp1Δ::KANMX6* cells may have been responsible for the mutant phenotypes we observed, rather than loss of Dbp1 protein.

**Figure 2.**
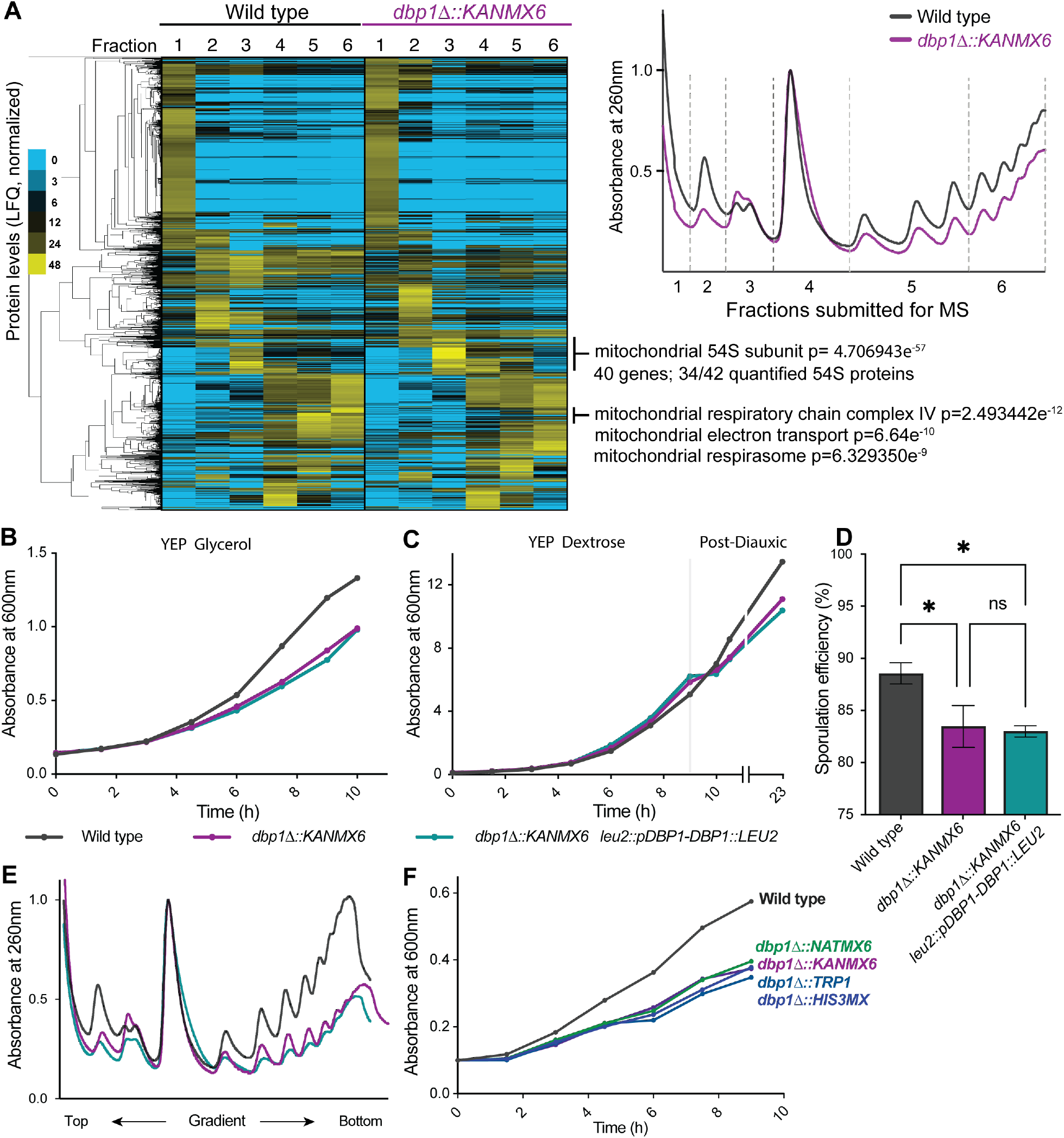
Resistance cassette-driven mis-regulation of *MRP51* causes widespread phenotypic consequences. (A) Label free mass spectrometry of fractionated polysomes from wild-type and *dbp1°::KANMX6* cells during meiosis. Note decreased polysomes, accumulation of mitochondrial 54S subunit (fraction 3), and a decrease in cytoplasmic 40S peak (fraction 2) for *dbp1°::KANMX6* cells. Two biological replicates were analyzed, one is shown. Replicates and their agreement by spearman correlation are shown in figure supplement. For full data see Figure 2 – Source Data 1. (B-E). Analysis of wild type, *dbp1°::KANMX6*, and *dbp1°::KANMX6 leu2::pDBP1-DBP1::LEU2* under conditions where *dbp1°::KANMX6* phenotypes were observed. (B) Growth curves in the non-fermentable carbon source glycerol, and ethanol (C: post-diauxic). (D) 24h sporulation efficiency counts, data shown is the average of 3 biological replicates with error bars showing SEM. Statistical significance was determined by a two-way ANOVA adjusted for multiple comparisons with Tukey’s multiple comparison test. (E) Polysome profiles from meiotic cells. Two biological replicates were analyzed, one representative trace is shown. (F) Growth curves of wild type compared to mutant strains where *DBP1* was replaced by varied resistance cassettes in the non-fermentable carbon source glycerol.

Disruption of mitochondrial translation, and ultimately function, would be expected to lead to global cellular fitness defects. Based on the apparent mitochondrial defects observed in the mass spectrometry data, we hypothesized that *dbp1Δ::KANMX6* cells would grow poorly in conditions where elevated mitochondrial function is required. Consistently, *dbp1Δ::KANMX6* cells exhibited severe growth defects in media containing only non-fermentable carbon sources, such as glycerol or acetate (Figure 2—figure supplement 1D). Additionally, growth defects were evident in *dbp1Δ::KANMX6* cells grown in rich media (YEP dextrose or sucrose) during post-diauxic growth stages, after saturated cultures have exhausted fermentable carbon source availability and begin to utilize ethanol through respiration (Figure 2—figure supplement 1E). To determine whether these phenotypes were due to cassette-induced *MRP51* mis-regulation, and not caused by loss of Dbp1 protein, we tested whether they could be rescued by exogenous expression of Dbp1. We expressed Dbp1 at endogenous levels in the *dbp1Δ::KANMX6* cells by integrating a single copy of the *DBP1* ORF and all its regulatory sequences at the *LEU2* locus. All *dbp1Δ::KANMX6* growth defects observed in conditions requiring respiratory growth remained similar with and without Dbp1 (Figure 2B-C), confirming the respiratory growth defects observed in *dbp1Δ::KANMX6* cells result from cassette insertion rather than lack of Dbp1 protein.

We next assessed whether our other observed phenotypes in *dbp1Δ::KANMX6* cells were also due to off-target effects, rather than loss of Dbp1 protein. Indeed, the sporulation defect observed in *dbp1Δ::KANMX6* cells was not rescued by Dbp1 (Figure 2D), which we now believe to result from the dependency of meiotic cells on respiratory function^48^. Finally, the reduced cytoplasmic 40S signal and diminished polysome levels representing lower translation rates in meiotic *dbp1Δ::KANMX6* cells was also not rescued by Dbp1 (Figure 2A, Figure 2—figure supplement 1A). Instead, we believe the bulk cytoplasmic translation phenotypes may reflect a cellular response to mitochondrial dysfunction or reduced mitochondrial translation rates, both of which have been shown to interact with the TOR pathway thus influencing global translation^49–52^. We conclude that resistance cassette insertion at the *DBP1* locus is sufficient to cause widespread cellular phenotypes not associated with loss of *DBP1* function. These data raised concerns as they confirm that resistance cassette-driven cis effects can confound the interpretation of phenotypes in mutants created in this manner. While these results were disappointing regarding our study of the function of Dbp1 helicase, they highlighted this locus as a useful context to dissect the mechanism behind, and ideally fix, resistance-cassette-related side effects.

To assess whether cassette-driven off-target effects were common to a specific feature of the KANMX6 and HYGMX4 selection cassettes, we replaced *DBP1* ORF in wild-type cells with three additional cassettes. Among the cassettes tested, we varied the promoter and terminator sequences used to express the resistance ORF, as well as the identity of the resistance ORF. Insertion of every resistance cassette tested lead to a similar growth defect in a non-fermentable carbon source (Figure 2F), a phenotype which was not dependent on Dbp1 (Figure 2B). We concluded that the cis effects exhibited with *DBP1* replacement were not specific to any one cassette and instead are a consequence of the resistance-mediated gene replacement approach.

**Figure 2—figure supplement 1.**
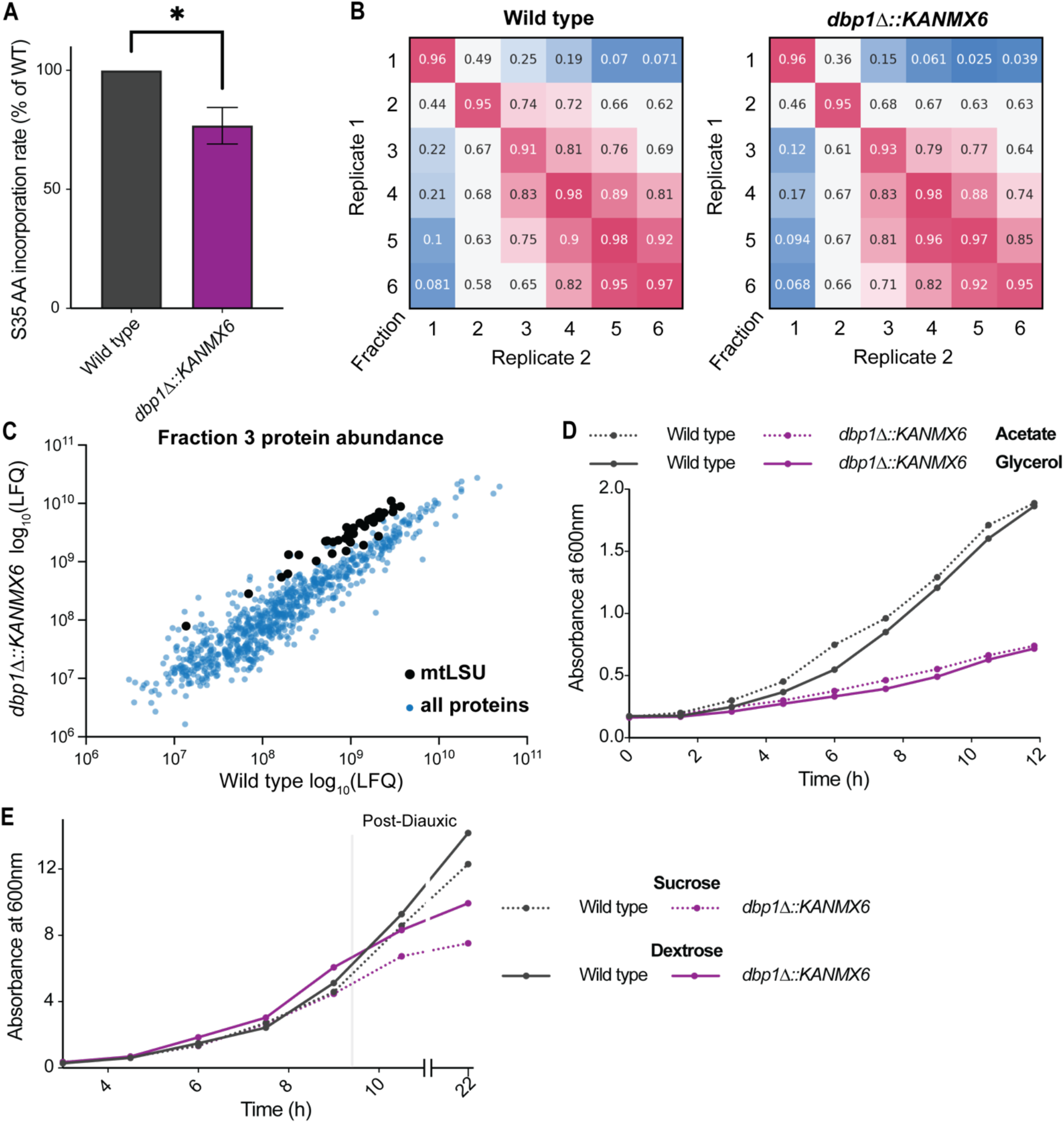
*DBP1* ORF replacement with KANMX6 causes diverse phenotypes resulting from mitochondrial dysfunction. (A) S35 amino acid incorporation of *dbp1°::KANMX6* and wild-type cells during meiosis. Statistical significance determined by an ordinary one-way ANOVA (p value <0.05). Data is average of three biological replicates; error bars show SEM. (B) Spearman correlation of biological replicates from label free quantification (LFQ) of fractionated polysome mass spectrometry experiment. (C) Normalized LFQ of all proteins quantified in fraction 3 shown as average of both replicates. (D-E) Growth curves of wild-type and *dbp1°::KANMX6* cells in non-fermentable carbon sources (D) and fermentable (E: before diauxic shift) and non-fermentable (E: post diauxic shift) carbon sources.

### Resistance cassette-induced aberrant transcription events are not specific to the *DBP1* locus

Because of the dramatic off-target effects observed with cassette replacement of *DBP1*, we wanted to determine if cassette insertion could drive aberrant transcription and neighboring gene misregulation at other loci. To assess this, we observed mRNA-seq reads over genomic intervals surrounding replaced ORFs, using previously published mRNA-seq and ribosome profiling data from our lab^47^ as well as an unpublished mRNA-seq dataset, also from our lab. All strains were made by replacing ORFs with either the KANMX6 or NATMX6 resistance cassettes and samples were collected during mitotic exponential growth in rich media. We found aberrant transcripts stemming from cassette-mediated gene replacements at 5 of these 19 additional observed loci (Figure 3 A-E). For each of these 5 new examples, the aberrant transcript appeared to be driven from the 5’ end of the resistance cassette, like in the case of *dbp1Δ::KANMX6* (Figure 1E, Figure 3 A-E). For 4 of these new cases in which aberrant transcription resulted from a cassette-based deletion, the adjacent gene was positioned in the same orientation as the aberrant TSS thus it would be possible for synthetic LUTI-based repression to occur as was observed for *MRP51*. However, unlike the strong mis-regulation of *MRP51* seen with *DBP1* ORF replacement, these aberrant transcription events had no obvious effect on the translation of the adjacent genes, consistent with the lack of apparent uORF translation in the extended 5’ mRNA regions (Figure 3A-D). Although, we also note that for 3 of these 4 cases, expression of the adjacent ORF was low, even in wild-type cells, and thus misregulation of these genes may be difficult to detect under these conditions compared to what we observed for *MRP51* (Figure 3B-D). For the last case, in which *UPF1* was replaced with the NATMX6 cassette, an abundant aberrant transcript was observed. However, the adjacent *ISF1* gene was oriented antisense to this transcript and there was no evidence of its transcription with or without NATMX6 insertion at *UPF1* (Figure 3E); hence, there was no baseline expression to disrupt. Together, these data suggest that cassette-induced-aberrant transcription events are not a phenomenon unique to the *DBP1*/*MRP51* genomic context, but that such severe phenotypic consequences may be more rare.

**Figure 3.**
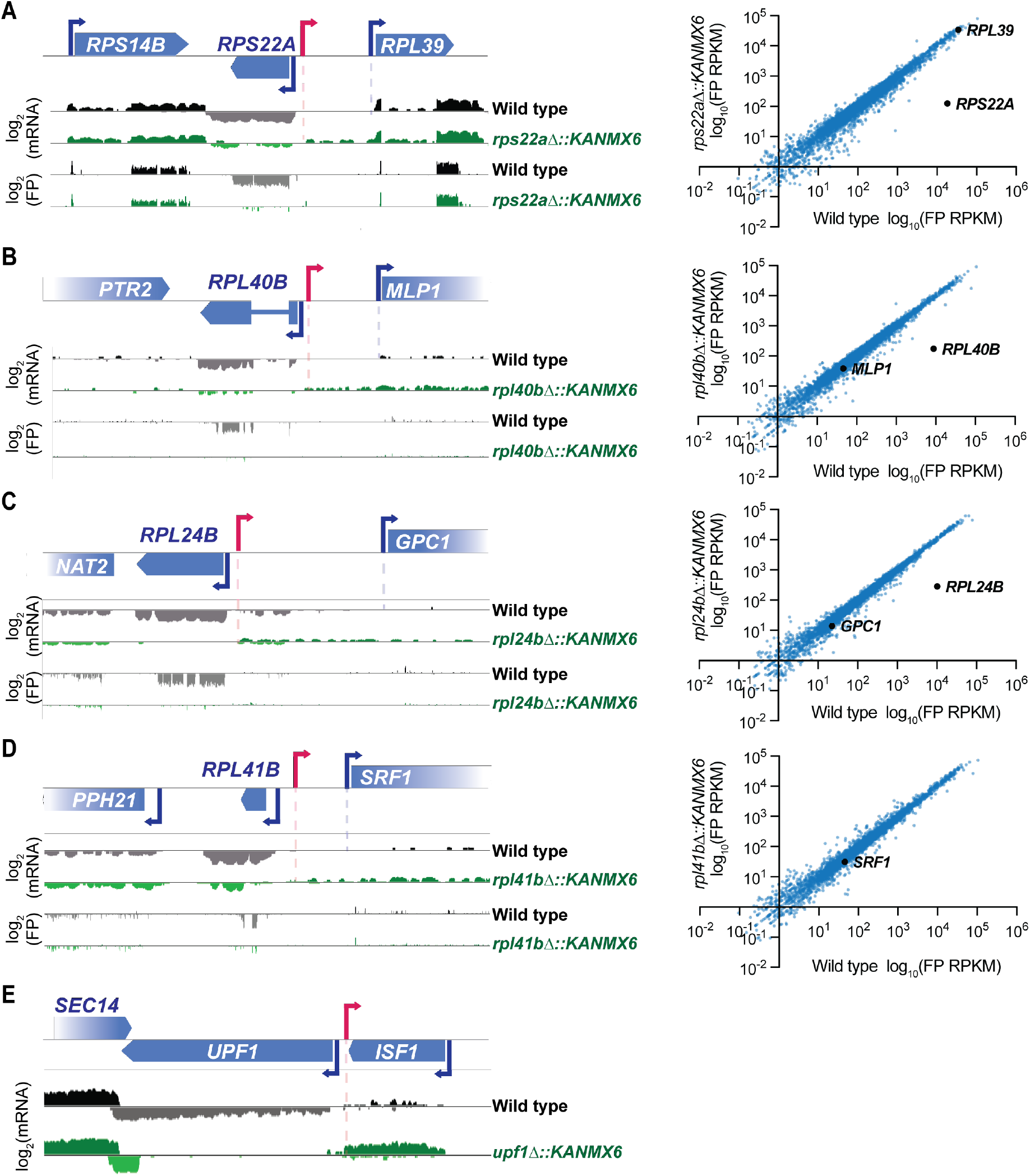
Resistance cassette-induced aberrant transcripts are not specific to the *DBP1* locus. (A-D) 18 ribosomal protein deletions generated by ORF replacement with KANMX6 from Cheng et al., 2018^47^, were observed for aberrant transcription events. Left, ribosome profiling footprint (FP) and mRNA-seq reads aligned to the ORF-replaced locus for each strain. Right, FP RPKM for every ORF expressed in the mutant and wild-type strains. Data was collected from vegetative exponentially growing cells. (D) Note: *RPL41B* is not quantified in translation graph because the ORF is only 25 amino acids long. (E) mRNA-seq reads aligned to the replaced locus in a *upf1°::NATMX6* mutant. Data was collected from vegetative exponentially growing cells. No matched ribosome profiling data was available for this study. See Figure 3 – Source Data 1.

The finding that deleting an ORF by replacement with a resistance cassette can cause mis-regulation of adjacent genes and potent off-target effects may seem to motivate the use of marker-less gene editing strategies for future studies. However, when we created a marker-less deletion of the *DBP1* ORF using CRISPR/Cas9 editing^7^ we observed surprising effects on the expression of surrounding genes as well. “Seamless” deletion of the *DBP1* ORF by this strategy led to the production of a transcript containing only the 5’ and 3’ UTRs of *DBP1*. Expression of this mutant transcript led to the aberrant translation of a dubious ORF contained within the *DBP1* 3’ UTR, not translated in wild type (Figure 3—figure supplement 1). These data, combined with the greater versatility of selection-marked mutations, highlight the need for improved modes of generating resistance marked mutations.

**Figure 3—figure supplement 1.**
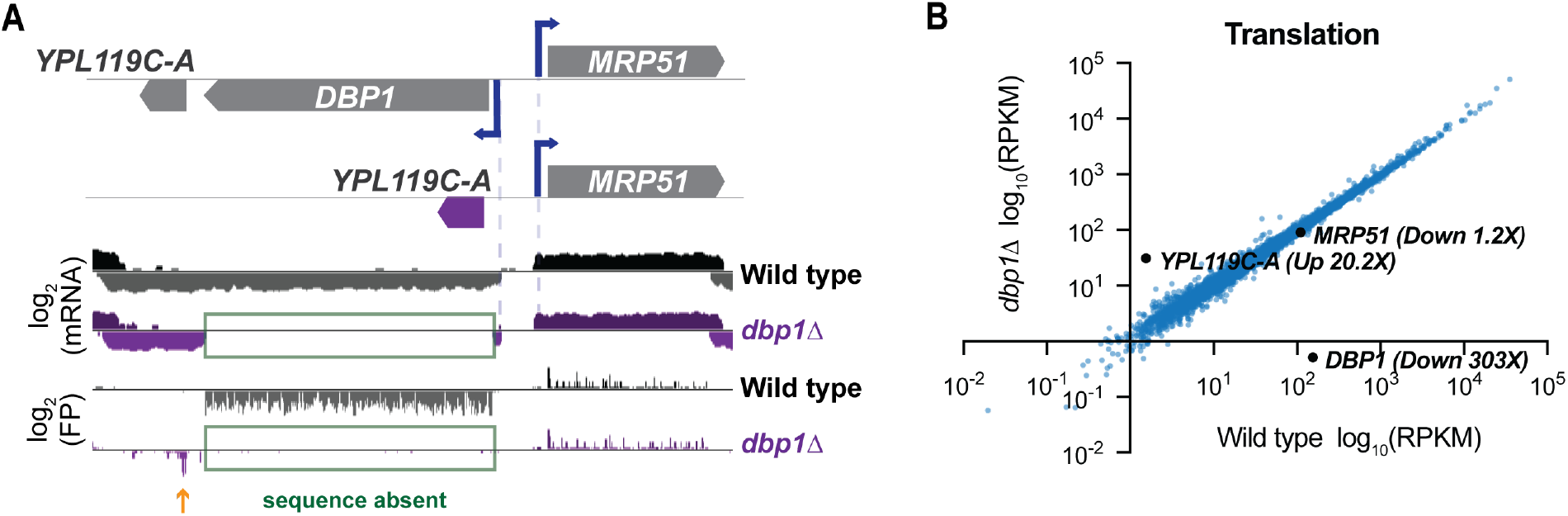
Cas9-mediated “seamless” deletion of *DBP1* ORF causes aberrant translation of an ORF within the *dbp1°* 3’UTR. (A) Ribosome profiling footprint (FP) and mRNA-seq reads mapped to the *DBP1* locus in wild-type and *dbp1°* (ORF removed with no resistance marker inserted) cells. Samples were collected during meiosis. (B) Translation (FP RPKM) for all genes expressed in *dbp1°* and wild-type cells during meiosis. See Figure 3—figure supplement 1-Source Data 1 for all quantified genes.

### Insulated resistance cassettes prevent the mis-regulation of insertion adjacent genes

Because bi-directional transcription has been shown to occur from many promoters^8,11,53^, and all observed aberrant transcripts came from the end of the cassette containing the resistance ORF promoter, we thought this promoter was the most likely cause of the aberrant transcription events. Thus, to circumvent these events from disrupting nearby genes, we placed strong transcription terminator sequences flanking both ends of the resistance cassette^54^. We placed a copy of the *DEG1* terminator 5’ to the *TEF1* promoter, and either the same *DEG1* terminator or the *CYC1* terminator 3’ to the *TEF1* terminator within the KANMX6 cassette (Figure 4A). We named these plasmids “KANMX6-ins” and used them to replace the *DBP1* ORF using the same primers and insertion location as before. Both *dbp1Δ::KANMX6-ins* strains grew at wild-type rates in the non-fermentable carbon source glycerol, in contrast to cells housing the original cassette replacement (Figure 4B). Consistent with this finding, levels of Mrp51 protein in the *dbp1Δ::KANMX6-ins* strains were similar to those in wild-type cells (Figure 4C, D). Additionally, 5’ rapid amplification of cDNA ends (5’RACE) confirmed that the *MRP51* transcript produced in the *dbp1Δ::KANMX6-ins* strains was the same as that seen in wild-type cells, as opposed to the synthetic *MRP51* LUTI in the initial cassette-inserted strains (Figure 4E—figure supplement 1A-B). We were pleased that this simple change to the resistance cassette structure prevented the neighboring gene mis-regulation and resulting systemic phenotypes seen previously with *DBP1* ORF replacement. These data suggest that this strategy may prove useful to prevent aberrant transcription and neighboring gene misregulation in future studies. We thus created insulated versions of cassettes housing three additional commonly used selection markers *TRP1, HIS5* (*S. Pombe HIS5*; which complements *his3*), and *NAT* (nourseothricin resistance) into the classic “MX6” backbone^1-3^. We believe that use of these insulated cassettes in place of the traditional set should drastically reduce off-target effects from future genetic studies.

**Figure 4.**
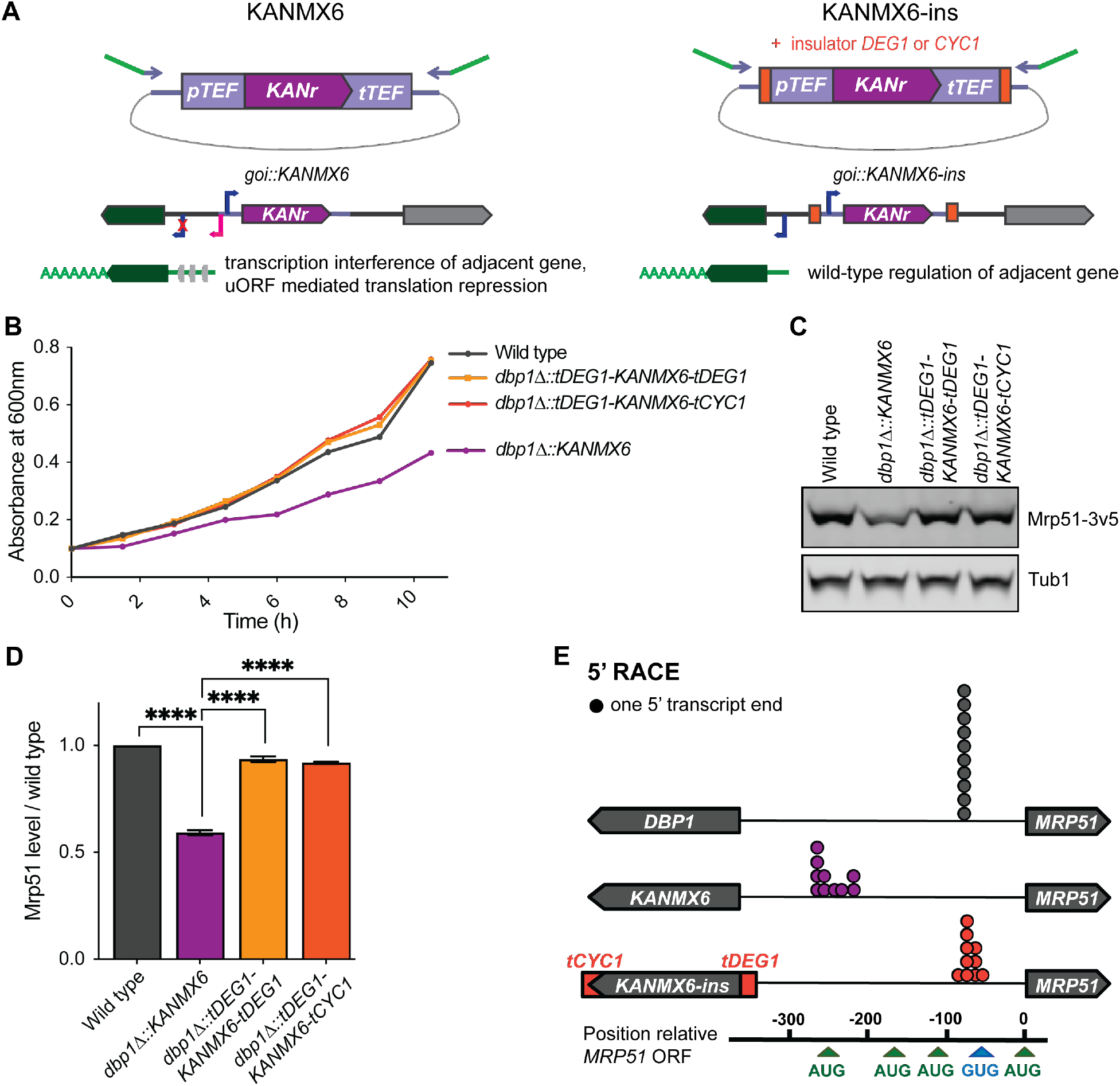
Insulated resistance cassettes prevent cassette-induced mis-regulation of adjacent genes. (A) Design of “insulated” KANMX6-ins cassettes and proposed model. The *DEG1* transcription terminator sequence was inserted 5’ of the *TEF1* promoter and either the *DEG1* or *CYC1* terminator was inserted 3’ of the *TEF1* terminator in pFA6a-KANMX6^1^. (B) Growth of *dbp1°::KANMX6* and *dbp1°::KANMX6-ins* strains compared to wild type in YEP-glycerol. (C) Western blots for Mrp51-3v5 levels in *dbp1°::KANMX6* and *dbp1°::KANMX6-ins* cells compared to wild type during mitotic exponential growth in rich media. For full blot scans see Figure 4 – Source Data 1. (D) Quantification of western blots in (C) with Mrp51 normalized to Tub1 (alpha tubulin). Data represent average of two biological replicates with error bars showing range. Statistical significance was determined by a one-way ANOVA adjusted for multiple comparisons using Tukey’s multiple comparisons test, P values less than .005 are shown. (E) 5’ rapid amplification of cDNA ends (RACE) demonstrates the 5’ ends of transcripts produced in wild type, *dbp1°::KANMX6, and dbp1°::KANMX6-ins* strains. Exact 5’ mRNA end sequences listed in figure supplement.

**Figure 4—figure supplement 1.**
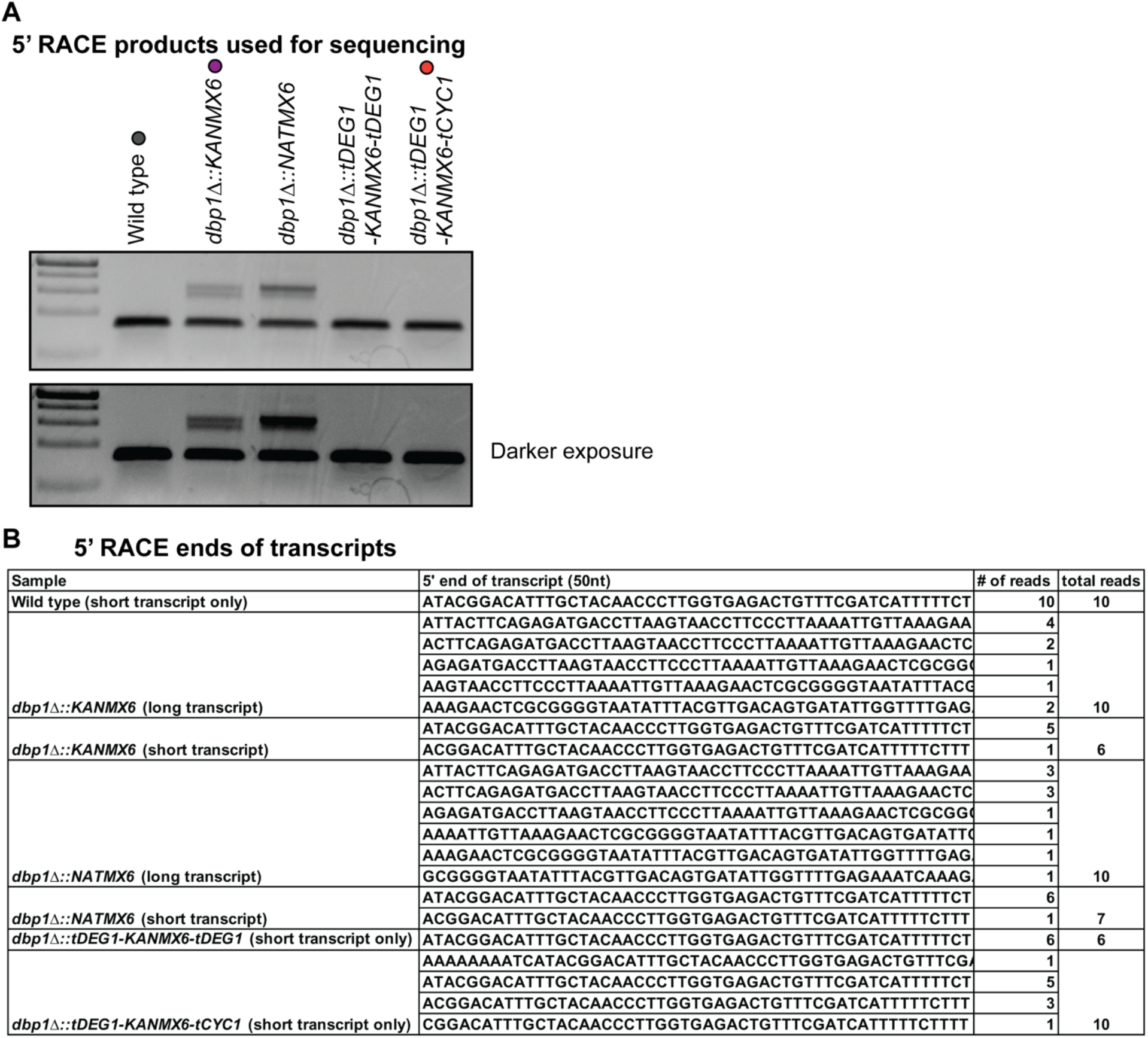
5’ RACE confirms the presence of wild-type 5’ end of *MRP51* transcript when *DBP1* ORF is replaced with KANMX6-ins cassettes. (A) PCR amplified 5’ ends of cDNA from wild type, *dbp1°::KANMX6, dbp1°::NATMX6, dbp1°::tDEG1-KANMX6-tDEG1*, and *dbp1°::tDEG1-KANMX6-tCYC1*. Colored dots represent the color each sample is depicted with in Figure 4E. Bottom image is darker exposure of the same gel to demonstrate lack of *MRP51* LUTI in insulated cassette samples. See Figure 4 – figure supplement 1 – Source Data 1 for uncropped images of gels. (B) The exact 5’ ends of transcripts sequenced to identify the start site of each transcript as well as the number of reads sequenced at each position.

## Discussion

In this study, we provide evidence that resistance-cassette-mediated genome engineering in budding yeast can result in overlooked and potent off-target effects. Dissection of the mechanism behind these off-target effects at the *DBP1*/*MRP51* loci demonstrate their potential to ablate expression of neighboring genes through use of cassette-induced alternative TSSs and production of repressive LUTIs. This regulation, integrates transcription interference and translation control, and occurs despite the ORF and normal regulatory sequences for the affected adjacent gene remaining intact at the genomic level^20,30,31^. This, combined with condition-specific manifestation of phenotypes caused by off-target effects, as was seen at the *DBP1*/*MRP51* locus, can make identification of the off-target effects difficult. Had we not generated mRNA-seq and ribosome profiling data and been aware of the features of naturally occurring LUTI regulation, we would have misinterpreted mutant phenotypes in the *dbp1Δ::KANMX6* cells as relevant to Dbp1 function. Additionally, the well characterized function of Mrp51 allowed us to test whether the cassette-mediated mis-regulation of *MRP51* was sufficient to cause cellular phenotypes. We believe cassette-induced mis-regulation of adjacent genes may be a more elusive problem in studies where genome-wide gene expression information is not generated and that this side effect can cause misinterpretation of mutant phenotypes.

How common is this unexpected effect? Confidence in the specificity of yeast genome-editing cassettes has motivated their widespread use and many landmark studies defining the functions of conserved proteins have been performed using this approach. It is routine for multiple loci to be disrupted within the same strain to enable analysis of genetic interactions, and global yeast deletion, hypomorphic, and epitope tag collections were created by use of this exact insertion cassette^55–61^. Our ability to identify cassette-driven aberrant transcripts in previous studies from ourselves and others demonstrate that this side effect can be induced with a variety of cassettes and at multiple loci. These data suggest this type of genome engineering artifact may be an overlooked and relevant factor in the interpretation of mutants from many published studies. Of the 20 selection-cassette-mediated loss-of-function mutants for which our lab had generated mRNA-seq data, 6 produced a clear aberrant transcript (Figure 1E, Figure 3 A-E). However, none of the 5 additional cases resulted in severe effects on translation of neighboring genes, as was seen with *DBP1/MRP51*.

We believe the dramatic shut down of Mrp51 synthesis resulted from the convergence of multiple genomic features allowing for maximal mis-regulation via the LUTI-based mechanism. These features included a gene antisense and 5’ to the cassette insertion, translated competitive AUG uORFs, and lack of an intervening transcriptional terminator between the replaced and adjacent ORF. The combination of all these features is not common to most loci but would be expected to be seen in many cases genome-wide. It is also worth noting that phenotypic consequences resulting from cassette mis-regulation of adjacent genes may be condition-specific, as was the case with *MRP51*. Exponentially growing *dbp1Δ::KANMX6* cells in rich media grow at wild-type rates despite having lower levels of Mrp51 protein, presumably a result of the reduced need for mitochondrial function under these conditions (Figure 2C: prior to the diauxic growth phase). Under conditions requiring elevated mitochondrial function, the *MRP51* mis-regulation becomes limiting and widespread phenotypes occur (Figure 2A-F, Figure 2— figure supplement F-G). The cassette-induced-aberrant transcription events reported here that did not have evident mis-regulation of adjacent genes under the single condition we assayed may cause further adjacent gene disruption and phenotypic consequences under conditions not tested in this study.

While we focused on the potential for integrated transcription and translation LUTI-based neighboring-gene disruption in this study, mis-regulation of adjacent genes by cassette-driven transcription interference alone may happen more frequently. This mis-regulation can occur regardless of the adjacent gene’s direction in relation to the cassette and does not require the transcript to be continuous over the adjacent TSS. For example, in haploid cells induction into meiosis is blocked by the expression of two separate non-coding RNAs that repress an antisense mRNA (*RME2*/*IME4*) or a downstream sense RNA (*IRT1*/*IME1*) via transcription interference. Transcription interference between loci can be effective over a large distance, like the ∼1800 nt span over which transcription of the repressive *IRT1* long noncoding RNA represses *IME1*^22^. Considering the number of factors that can affect whether a gene adjacent to a resistance cassette could be mis-regulated, we believe it is hard to predict when and where the side effect reported here would result in misregulation severe enough to mislead studies. Two recent studies analyzing the non-essential yeast deletion collection show clear signs of the cassette driven off-target effects reported here and indicate that this effect may be more common than our analyses suggest. One study found that one of the most significant factors determining whether a specific protein’s abundance would change following deletion of a gene encoding another protein was whether the genes encoding the protein pair were genomic neighbors^62^. Another study, which analyzed genetic interaction data, determined that neighbor gene effects of unknown mechanism appeared to account for non-specific phenotypes in ∼ 10% of deletion strains^63^.

What is driving aberrant transcription from selection cassette insertion? It has been recently shown that most promoters in yeast are bidirectional and that resulting antisense transcripts are often not evident due to their targeting by mRNA decay mechanisms^8,12,53^. We believe the aberrant transcripts in our study arise from bidirectional transcription stemming from promoters within the cassette used to express resistance ORFs. Instability of such antisense transcripts in RNA-decay-proficient backgrounds may explain why we do not observe aberrant transcripts driven by cassettes in 14 of the 20 cases we analyzed. Thus, we may be underestimating the frequency by which adjacent genes are affected by cassette-mediated transcription interference. Alternatively, it is possible that the chromatin context of cassette insertion controls the bidirectional activity of the cassette promoters. Studies using resistance cassette generated mutants in strain backgrounds incompetent for mRNA decay will be informative in deciphering between these possibilities. Interestingly, the *upf1Δ::NATMX6* antisense transcript produced appeared to be more abundant than many of our other examples (Figure 3E compare to A-D). Upf1’s role as an RNA helicase that drives degradation of mRNAs through the non-sense mediated decay pathway^64,65^ could be a relevant factor to investigate in future studies.

A major challenge presented by the cassette-mediated off-target effects we found in this study is that they are difficult to detect. PCR checking, sequencing, and backcrossing of cassette-inserted strains are common approaches to identify or prevent potential non-specific mutations and phenotypes, but none of these strategies can prevent or detect cassette-driven aberrant transcription. No unexpected mutations exist in these cases and even mRNA measurements of neighboring genes are not diagnostic unless an approach is designed to assess all transcript isoforms from a given locus. We find that measurement of translation or protein levels from neighboring genes can be diagnostic of this undesirable side-effect, but ribosome profiling is not a routine experiment for analysis of most mutants, and tagging of many proteins does not retain their normal function and regulation. Additionally, as many genes are expressed in a condition-specific manner, comparison of the adjacent gene’s protein levels in a mutant as compared to wild type may give false confidence unless the proper conditions are chosen. Here, for 3 of the 6 mutants with observable aberrant transcripts that could disrupt adjacent genes via LUTI-based regulation, those genes were lowly expressed, even in wild-type cells, under the conditions surveyed.

Despite this complexity, use of complementation-based rescue experiments— namely single copy insertion of the deleted gene with its entire endogenous regulatory region at another locus— allowed us to clearly identify the unexpected regulation in the case of *DBP1* disruption. This type of approach should be more commonly used to confirm mutant phenotypes, even in “seamless” deletes, but may be complex or unfeasible in cases in which cassette insertion is used for epitope tagging or experiments in which genetic interactions between several genes are being investigated.

We suggest the future use of “insulated” resistance cassettes to preventing off-target effects and misinterpretation of mutant phenotypes in future genome editing experiments. Precision Cas9-based ORF deletion is a also powerful approach to genome editing, but can also result in changes to the chromatin context of an adjacent gene, potentially resulting in LUTI-based repression^13,7^. For the case of a Cas9-generated marker-less deletion of the *DBP1* ORF, we observed translation of a previously untranslated ORF within the mutant transcript containing only the *DBP1* 5’ and 3’ UTR (Figure 3—figure supplement 1). One could imagine that in cases like these, translation of novel ORFs could also cause misleading neomorphic mutant phenotypes not linked to the deleted ORFs loss of function. While our study was limited to yeast, all factors behind the cassette-induced off-target effects identified here are conserved to higher eukaryotes and we believe genome-editing-driven LUTI-based regulation is likely to occur in other organisms. Our data highlight this as a previously overlooked side-effect of genome-editing which should be considered in the analysis of edited cells. They also emphasize the need for proper controls to establish causality of observed phenotypes to the intended mutation, even when using precision genome engineering approaches in yeast and other organisms.

## Acknowledgments

We thank N. Ingolia, E. Ünal, and current members of the Brar and Ünal labs for their comments on this manuscript. This work was supported by National Institutes of Health [1R35GM134886] funding to G.A.B. E.N.P was funded by an NSF predoctoral Fellowship.

## Materials and Methods

### Strains

All strains were derived from the Saccharomyces cerevisiae SK1 background^44^. Detailed strain genotypes can be found in supplemental table 1. Strains were diploid *MATa/alpha* unless otherwise specified as haploid (MATa). Selection-mediated loss of function mutants were created by transforming wild-type haploids with a linear PCR fragment encoding a selection cassette that would replace the ORF of interest via homologous DNA repair mechanisms^(1-5)^. Sequences directly upstream and downstream of each replaced ORF were always left intact. Genotypes for all strains were confirmed using PCR and mutants were backcrossed to a wild-type strain of the opposite mating type to ensure clearance of possible background mutations from transformation. The marker-less *dbp1Δ* was created by transforming a wild-type haploid with a plasmid encoding Cas9 and a sgRNA targeting the *DBP1* ORF, alongside a linear repair fragment that removes the *DBP1* ORF while preserving the upstream and downstream adjacent sequence. Strains containing the Mrp51-3v5 allele were created by transforming wild type or the selection replaced *dbp1Δ* strain of interest with a plasmid encoding Cas9 and a sgRNA targeting *MRP51*, alongside a linear repair template that c terminally tags Mrp51 with the 3v5 sequence and introduces a synonomous mutation in the guide target sequence to prevent re-editing. Strains expressing exogenous Dbp1 from the *LEU2* locus were created by cloning the *DBP1* ORF as well as ∼1kb of upstream and downstream regulatory sequence into a single integration plasmid targeted to *LEU2* This plasmid was linearized by digestion with SwaI nuclease (NEB, Ipswich, MA) and transformed into the *dbp1Δ::KANMX6* strain.

### Yeast Growth Conditions

Yeast were grown for experiments as in (Powers et al., 2021). Briefly, strains were thawed on YPG plates from glycerol stocks overnight, then grown on YPD plates for a day before use in experiments. All growth experiments were carried out at 30°C using YEP media supplemented with 2% w/v of the carbon source, except for glycerol which was used at 3% v/v. All growth experiments shown were repeated at least twice with one representative growth curve shown for each. Synchronous sporulating cells were prepared by first growing cells in YEP 2% dextrose for 24 hours at room temperature then diluted into BYTA at OD_600_ = .25 and grown overnight at 30°C. Next, cells were pelleted, washed with water, and resuspended in sporulation (SPO) medium: either (0.5% KAc [pH = 7.0] supplemented with 0.02% raffinose) for Figure 1 ribosome profiling and mRNA-seq experiments, or rich SPO medium (2% KAc [pH = 7.0] supplemented with 40 mg/L adenine, 40 mg/L uracil, 20 mg/L histidine, 20 mg/L leucine, 20 mg/L tryptophan for sporulation efficiency, polysome experiments, and ribosome profiling in Figure 3—figure supplement 1. Sporulation efficiency was counted at 24 hours after induction into SPO media and 200 cells were counted per sample for 3 independent biological replicates. Samples were blinded prior to counting.

### Ribosome profiling and mRNA-seq

Meiotic yeast cells were harvested after 4 or 6 hours in sporulation medium (Figure 1 data: 4 hours; Figure 3—figure supplement 1: 6 hours), by brief treatment with 100uM then flash freezing. They were subjected to ribosome profiling and mRNA-seq as described in^37^. Reads per ORF were determined following mapping of all reads to the SK1 genome. RPKM values for ribosome footprints (FP) and mRNA reads were calculated by dividing the number of raw reads per gene by the total number of million mapped reads per sample and by the length in kilobases of each gene. Translation efficiency (TE) was calculated by dividing the FP RPKM / mRNA RPKM for each gene. Data visualization of genome tracts was made using MochiView^66^.

### Immunoblotting

Trichloroacetic acid (TCA) extractions were performed to collect total protein from samples as described in (Chen 2017)^30^. Briefly, ∼2.5OD_600_ of yeast were treated with 5% TCA at 4°C overnight, washed with acetone, dried then lysed in lysis buffer (50 mM Tris–HCl [pH 7.5], 1 mM EDTA, 2.75 mM DTT, protease inhibitor cocktail (cOmplete EDTA-free, *Roche*) for 5 minutes on a Mini-Beadbeater-96 (*Biospec Products*). Next, 3x SDS sample buffer (187.5 mM Tris [pH 6.8], 6% ß-mercaptoethanol, 30% glycerol, 9% SDS, 0.05% bromophenol blue) was added to 1X and the cell lysate was boiled for 5 min. Proteins were run on 4-12% Bis-Tris Bolt gels (Thermo Fischer) then transferred to .45uM nitrocellulose membranes using the 30 minute mixed molecular weight protocol on the Trans-Blot Turbo System from Bio-rad. Membranes were blocked with Intercept (PBS) blocking buffer (*LI-COR Biosciences, Lincoln, NE)* at room temperature then incubated overnight at 4°C with mouse anti-v5 (1:1000, Invitrogen, RRID:AB_2556564), and rat anti-tubulin alpha (1:10,000, Serotec, RRID:AB_325005). Membranes were washed in PBS-.08% tween then incubated with an anti-mouse secondary antibody conjugated to IR Dye 800 (RRID:AB_621842) at a 1:15,000 dilution, and either an anti-rat secondary antibody conjugated to IR Dye 680 (RRID:AB_10956590) at a 1:15,000 dilution at room temperature for 1-2h (LI-COR Biosciences). Immunoblot images were generated and quantified using the Odyssey system (*LI-COR Biosciences*).

### Polysome Analysis

Cells were treated with 100uM cycloheximide for 30s then filter collected, flash frozen, lysed, and prepared for sucrose gradient centrifugation of as in^67^, with the following exceptions. Samples were thawed just prior to their loading on sucrose gradients, and SUPERase·In was added as for “mock digested samples”. Also, the polysome lysis buffer used was modified to contain protease and phosphatase inhibitors and is as follows 20mM Tris pH8, 140mM KCl, 1.5mM MgCl2, 100ug/ml cycloheximide, 1% Triton X-100, 2ug/ml Aprotinin, 10ug/ml Leupeptin, 1mM PMSF, 1:100 PIC2, 1:100 PIC3. Lysates were loaded onto 7-47% sucrose gradients and spun in a SW 41 Ti rotor for 3 hours at 35,000rcf at 4°C. Following the spin, samples were kept at 4°C and immediately collected using the Gradient Master gradient station from BioComp while monitoring 260nm absorbance with the BioRad EM-1 Economonitor. Samples were fractionated as shown in Figure 2, flash frozen, and submitted for mass spectrometry analysis.

### Polysome fraction processing for LC-MS/MS measurements

Proteins were precipitated and desalted using the SP3 method, as described in (Hughes et al., 2018)^68^. 50% from each polysome fraction (volume) were processed. Disulfide bonds were reduced with 5mM dithiothreitol and cysteines were subsequently alkylated with 10mM iodoacetamide. Proteins were precipitated on 0.5 µg/µL speedBead magnetic carboxylated modified beads (1:1 mix of hydrophobic and hydrophilic beads, cat# 6515215050250, 45152105050250, GE) by addition of 100% ethanol in a 1:1 (vol:vol) sample:ethanol ratio followed by 15 min incubation at 25°C, 1000 rpm. Protein-bound beads were washed in 80% ethanol and proteins were digested off the beads by addition of 0.8ug sequencing grade modified trypsin (Promega) in 100mM Ammonium bicarbonate, incubated 16 hrs at 25°C, 600 rpm. Beads were removed and the resulting tryptic peptides evaporated to dryness in a vacuum concentrator. Dried peptides were further desalted by another round of SP3 precipitation; Peptides were reconstituted in 200ul of 95% Acetonitrile (ACN), followed by addition of bead-mix to a final concentration of 0.5 µg/µL and incubated 15 min at 25°C, 1000 rpm. Beads were subsequently washed in 80% ethanol, peptides were eluted off the beads in 50 µL 2% DMSO and samples were evaporated to dryness in a vacuum concentrator. Dried peptides were then reconstituted in 3% ACN / 0.2% Formic acid to a final concentration of 0.5 µg/µL.

### LC-MS/MS analysis on a Q-Exactive HF

About 1 μg of total peptides were analyzed on a Waters M-Class UPLC using a 25cm Ionopticks Aurora column coupled to a benchtop Thermo Fisher Scientific Orbitrap Q Exactive HF mass spectrometer. Peptides were separated at a flow rate of 400 nL/min with a 190 min gradient, including sample loading and column equilibration times. Data was acquired in data dependent mode using Xcalibur software. MS1 Spectra were measured with a resolution of 120,000, an AGC target of 3e6 and a mass range from 300 to 1800m/z. Up to 12 MS2 spectra per duty cycle were triggered at a resolution of 15,000, an AGC target of 1e5, an isolation window of 1.6 m/z and a normalized collision energy of 27.

All raw data were analyzed with MaxQuant software version 1.6.10.43 ^69^ using a UniProt yeast database (release 2014_09, strain ATCC 204508 / S288c), and MS/MS searches were performed with the following parameters: Oxidation of methionine and protein N-terminal acetylation as variable modifications; carbamidomethylation as fixed modification; Trypsin/P as the digestion enzyme; precursor ion mass tolerances of 20 p.p.m. for the first search (used for nonlinear mass re-calibration) and 4.5 p.p.m. for the main search, and a fragment ion mass tolerance of 20 p.p.m. For identification, we applied a maximum FDR of 1% separately on protein and peptide level. “Match between the runs” was activated, as well as the “LFQ” normalization (at least two ratio counts were necessary to get an LFQ value). We required 1 or more unique/razor peptides for protein identification and a ratio count of 2 or more for label free protein quantification in each sample. This gave intensity values for a total of 2143 protein groups across both replicates. “LFQ” normalized values were used for all subsequent analyses.

LFQ normalized values were then clustered with Cluster 3.0^70^ and visualized with Java TreeView^71^. Proteins not quantified in 2 or more samples were excluded in clustering. GO enrichment was assessed on the clusters specified (June 2022) and compared to the background population of all proteins quantified in the experiment (Boyle EI, et al. 2004).

### Radioactive amino acid incorporation assays

Cells were transferred to sporulation (SPO) media and incubated at 30°C for 4 hours. To metabolically label the cells 5uL of EasyTag EXPRESS ^35^S protein labeling mix (PerkinElmer), was added to 10mL of SPO cultures and incubated with shaking for 10 minutes at 30°C. Protein was precipitated by addition of 100uL 100% trichloroacetic acid to 900uL SPO culture and incubated at 95°C with shaking. Samples were chilled on ice then pelleted and washed with cold 10% trichloroacetic acid, followed by a wash in cold 100% ethanol. Samples were resuspended in 5mL of Econo-Safe scintillation fluid (RPI). Scintillation was counted for 2 minutes and the ^35^S incorporation rates were derived from counts per minute normalized to cell density between samples and normalized to wild type.

### Design and testing of insulated selection cassettes

Based on the previous report that the Deg1 and Cyc1 terminators can be used as insulator sequences^72^, we amplified these terminators from wild-type yeast and placed them into the 5’ and 3’ ends of previously designed selection cassettes using gibson assembly just inside the primer amplification sites. We chose to conserve the primer amplification sites of these plasmids so that they would be useable with any primers designed to the previous system (pFA6a)^1^. Insulated plasmids were fully sequenced over cloning junctions and the entire region to be amplified and integrated into yeast during transformation. To test the ability of these insulators to prevent aberrant transcription at the *DBP1* locus, the KANMX6-ins cassettes were amplified and transformed into yeast exactly as had been done for the previous KANMX6 selection cassettes used in this study. When used to replace *DBP1* ORF, Mrp51-3v5 levels and 5’ RACE confirmed the efficacy of the KANMX6-ins cassettes in preventing aberrant LUTI regulation of *MRP51*. We then cloned 3 additional cassettes using varying selection (including *Saccharomces pombe* HIS5: complements *S*.*cerevisiae* HIS3, TRP1, and NATr) such that they can be used conveniently in combination as was done for the original toolkit. Descriptions of these plasmids can be found in supplemental table 1.

### 5’ RACE

5’ RACE was performed with total RNA extracted from samples using phenol chloroform precipitation and the GeneRacer™ Kit with SuperScript III™ RT and Topo TA Cloning™ Kit for Sequencing. For each sample, 4.8ug of total RNA was used. Random primers were used to reverse transcribe the de-capped cDNA library. The 5’ ends of the *MRP51* cDNA were then amplified using the GeneRacer™ 5’ primer and a reverse gene specific primer (5’ ACCGTCCAAGCAACTCTGCCAATGTC 3’) in a reaction with Platinum™ *Taq* High Fidelity DNA Polymerase. SnapGene was used to visualize and align sequenced cDNA ends to *MRP51* sequence.

## Materials availability

All newly created materials are available upon request. Newly created “Insulated” selection cassettes will also be deposited to Addgene and made publicly available.

## Data Availability

We are currently in the process of uploading all relevant newly created datasets to public databases. Repository information will be shared when available. Source data for all figures generated from newly created datasets is currently shared within the manuscript.

## Source Data Information

**Figure 1 – Source Data 1:**

RPKM values for mRNA-seq and ribosome profiling data for all genes quantified in the experiment shown in Figure 1 plots.

**Figure 1 – Source Data 2:**

Zipped folder containing uncropped tif image of western blot shown in Figure 1 with and without labels on the relevant samples and bands.

**Figure 2 – Source Data 1:**

Label free quantification (LFQ) of all proteins quantified in fractionated polysome experiment shown in Figure 2 and Figure 2 – figure supplement 1.

**Figure 3 – Source Data 1:**

Zipped folder containing .txt (wiggle files) used to generate Figure 3E genome browser track figure. *UPF1* ORF coordinates 415257 – 418173 on chr 13. 2KB upstream of *UPF1* and 1 KB downstream of *UPF1* are included.

**Figure 3 – figure supplement 1 – Source Data 1:**

RPKM values for all genes quantified in ribosome profiling experiment shown in Figure 3 – figure supplement 1.

**Figure 4 – Source Data 1:**

Zipped folder containing uncropped tif image of western blot shown in Figure 4 with and without labels on the relevant samples and bands.

**Figure 4 – figure supplement 1 – Source Data 1:**

Zipped folder containing uncropped tif image of agarose gel showing the amplified 5’ RACE products sequenced in Figure 4 and shown in Figure 4 – figure supplement 1.

